# Linear superposition of responses evoked by individual glottal pulses explain over 80% of the frequency following response to human speech in the macaque monkey

**DOI:** 10.1101/2021.09.06.459204

**Authors:** Tobias Teichert, G. Nike Gnanateja, Srivatsun Sadagopan, Bharath Chandrasekaran

## Abstract

The frequency-following response (FFR) is a scalp-recorded electrophysiological potential that closely follows the periodicity of complex sounds such as speech. It has been suggested that FFRs reflect the linear superposition of responses that are triggered by the glottal pulse in each cycle of the fundamental frequency (F0 responses) and sequentially propagate through auditory processing stages in brainstem, midbrain, and cortex. However, this conceptualization of the FFR is debated, and it remains unclear if and how well a simple linear superposition can capture the spectro-temporal complexity of FFRs that are generated within the highly recurrent and non-linear auditory system. To address this question, we used a deconvolution approach to compute the hypothetical F0 responses that best explain the FFRs in rhesus monkeys to human speech and click trains with time-varying pitch patterns. The linear superposition of F0 responses explained well over 90% of the variance of click train steady state FFRs and well over 80% of mandarin tone steady state FFRs. The F0 responses could be measured with high signal-to-noise ratio and featured several spectro-temporally and topographically distinct components that likely reflect the activation of brainstem (<5ms; 200-1000 Hz), midbrain (5-15 ms; 100-250 Hz) and cortex (15-35 ms; ~90 Hz). In summary, our results in the monkey support the notion that FFRs arise as the superposition of F0 responses by showing for the first time that they can capture the bulk of the variance and spectro-temporal complexity of FFRs to human speech with time-varying pitch. These findings identify F0 responses as a potential diagnostic tool that may be useful to reliably link altered FFRs in speech and language disorders to altered F0 responses and thus to specific latencies, frequency bands and ultimately processing stages.

## Background

The frequency-following response (**FFR**) is a scalp-recorded electrophysiological potential that closely follows the periodicity of complex sounds such as speech (Aiken and Picton, 2008; Chandrasekaran and Kraus, 2010; Skoe and Kraus, 2010). Initially thought to reflect activity arising mostly from the cochlear nucleus and inferior colliculus (Chandrasekaran and Kraus, 2010) current thinking assumes multiple sources distributed across brainstem, midbrain and cortex (Coffey et al., 2019). Altered FFRs have been established as an important biomarker in speech and learning disorders (Cunningham et al., 2001; Banai et al., 2005, 2009; Chandrasekaran et al., 2009; Russo et al., 2009; Anderson et al., 2010; Hornickel et al., 2012; Hornickel and Kraus, 2013). Given the current view of the FFR as a signal arising from widely distributed sources, there are many different potential anatomical substrates for pathologically altered FFRs. Despite important advances, it has remained challenging to map altered FFR features to altered processing in specific brain regions. As a result, the potential of the FFR to reveal insights into specific circuits at different auditory processing stages has not been fully unlocked.

For *‘classical’* auditory evoked onset responses, important information about the neural origin can be gleaned from their latency and topography. Depending on their latency, neural responses have been coarsely attributed to auditory brainstem (<10 ms), midbrain (10-50 ms) or cortex (>50 ms) (Alain and Winkler, 2012). Topography, i.e., the spatial distribution of electric or magnetic fields across the scalp, can then be analyzed using source modeling approaches to further narrow down the exact spatial location of the underlying neural generators. Recent work has shown that source modeling can also be leveraged to better understand the neural generators of the FFR (Gerken et al., 1975; Bidelman, 2015; Coffey et al., 2016; Gorina-Careta et al., 2021). However, because of its dependence on high channel-count EEG and/or MEG recordings, source modeling is often not feasible for clinical FFR data which is typically recorded with a 3-electrode montage.

An alternative approach can be derived from the hypothesis that FFRs reflect the linear superposition of responses to each glottal pulse (F0 response) that sequentially activates processing stages in brainstem, midbrain, and cortex (Gerken et al., 1975; Janssen et al., 1991; Dau, 2003; Bidelman, 2015; Teichert et al., 2020). Despite its theoretical relevance, the superposition hypothesis has not been subject to much empirical scrutiny (Bidelman, 2015). If the superposition hypothesis is accurate, FFRs would arise as the convolution of the F0 response with a series of Dirac pulses whose time and amplitude reflect the onset and intensity of each glottal pulse, or more generally, each F0 cycle. Furthermore, it should be possible to compute the underlying F0 responses by inverting the convolution operation. So-called ‘deconvolution’ approaches have successfully been used in a wide range of neuroscientific applications (Aquino et al., 2014; Teichert and Ferrera, 2015), including the closely related 40 Hz auditory steady state response (Bohórquez and Özdamar, 2008) and continuous speech (Maddox and Lee, 2018; Polonenko and Maddox, 2021). To date, however, deconvolution has never been used to recover the F0 response underlying FFRs to stimuli with time-varying pitch. Thus, it is unknown how well a linear superposition model can account for the considerable spectro-temporal complexity of FFRs, and how much of their variance it can capture. If the F0 responses indeed account for a substantial portion of the FFR, they would provide useful information about FFR generators. Specifically, the latencies at which F0 responses differ could provide information about the latency of the altered processing stages. If, however, F0 responses only account for a very small fraction of the FFR, the F0 responses would likely be of limited use for understanding the alterations of the FFR in disease.

The goal of the current study was to determine if F0 responses can help link altered FFRs to altered function in specific auditory processing stages. To that aim we addressed three main questions: (i) What percentage of the variance of FFRs can be explained by the linear superposition of F0 responses? (ii) How reliably can F0 responses be estimated? (iii) Can the latencies of F0 responses be linked to anatomically distinct processing stages? Experiments were performed in macaque monkey which are known to exhibit human-like FFRs (Steinschneider et al., 1998, 2003; Brugge et al., 2009; Fishman et al., 2013; Ayala et al., 2017; Gnanateja et al., 2021). In addition, this species will ultimately allow us to confirm the results of the deconvolution method by directly measuring FFRs using invasive recordings.

## Methods

### Subjects

Data reported here was collected from 2 adult male macaque monkeys (*Macaca mulatta*). All experiments were performed in accordance with the guidelines set by the U.S. Department of Health and Human Services (National Institutes of Health) for the care and use of laboratory animals. All methods were approved by the Institutional Animal Care and Use Committee at the University of Pittsburgh. The animals had previously been exposed to pure tone and click-stimuli in passive and active listening paradigms.

### Stimuli

Two types of stimuli were used: (a) synthesized Mandarin tones (**Fig 1**A) and (b) click train versions thereof (**Fig 1**B). **Mandarin tones**: The synthesized mandarin tones used the vowel /yi/ in the context of 4 distinct F0 patterns: T1 (high-level, F0 = 129 Hz), T2 (low-rising, F0 ranging from 109 to 133 Hz), T3 (low-dipping, F0 ranging from 89 to 111 Hz), and T4 (high-falling, F0 ranging from 140 to 92 Hz). Mandarin tones were synthesized based on the F0 patterns derived from natural male speech production(Xie et al., 2017). All stimuli had a sampling rate 96000 Hz and were 250 ms in duration. The stimuli were presented in both condensation and rarefaction polarities to minimize potential contamination of the neural responses by the stimulus artifact and pre-neural cochlear microphonics^7^. The stimuli were presented in a randomized manner, with a randomly selected inter-stimulus intervals between 300 and 500 ms. In each 40-minute long recording session, we presented 500 repetitions of each tone and polarity for a total of 4000 sweeps. **Click train stimuli**: From each of the four synthesized mandarin tone stimuli we prepared a click train version that consisted of trains of 0.1 ms long monophasic impulses. Timing and amplitude of the clicks in the click trains matched the timing and amplitude of the F0 cycles of the mandarin tone stimuli. The timing of the F0 cycles was operationalized as the time of the peak pressure (**Fig 1**C, second F0 cycle), the intensity was operationalized as twice the absolute amplitude of the peak activity. The rationale for choosing twice the amplitude was to account for the fact that speech sounds are modulated bi-directionally.

**Figure 1.**
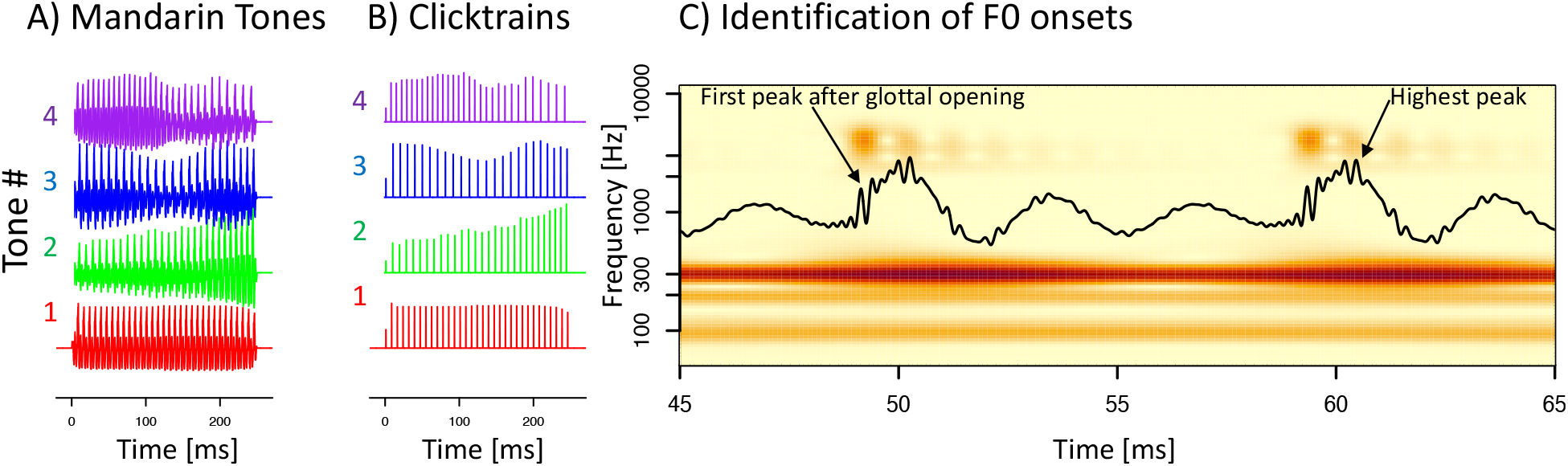
Stimuli. A) The 4 synthetic mandarin tone in the time domain. B) The corresponding click train stimuli. C) A snippet containing two F0 cycles of a mandarin tone stimulus in the time- (black line) and time-frequency domain (color). Timing of the clicks in the click train stimuli matched the time of the highest pressure peak (second F0 cycle). We subsequently defined the onset of an F0 cycle in as the first positive pressure peak that coincides with the first of several peaks of power in the third formant that follows the opening of the glottis (first F0 cycle).

### Experimental setup

All experiments were performed in a small (4’ × 4’ × 8’) sound-attenuating and electrically insulated recording booths (Eckel Noise Control Technology). The animal was positioned and head-fixed in custom-made primate chairs (Scientific Design). Neural signals were recorded at a sampling rate of 30kHz with a 256-channel digital amplifier system (*RHD2000, Intan*).

Experimental control was handled by a windows PC running an in-house modified version of the Matlab software-package *monkeylogic*. Sound files were generated prior to the experiments and presented by a sub-routine of the Matlab package *Psychtoolbox*. The sound-files were presented using the right audio-channel of a high-definition stereo PCI sound card (M-192 from M-Audiophile) operating at a sampling rate of 96 kHz and 24 bit resolution. The analog audio-signal was then amplified by a 300 Watt amplifier (QSC GX3). The amplified electric signals were converted to sound waves using a single element 4 inch full-range driver (Tang Band W4-1879) located 20 cm in front of the animals.

To determine sound onset with high accuracy, a trigger signal was routed through the unused left audio channel of the sound card directly to one of the analog inputs of the recording system. Thus, sound onset could be determined at a level of accuracy that was limited only by the sampling frequency of the recording device (30kHz: corresponding to 33 μs).

### Cranial EEG

EEG activity was recorded from 33 EEG electrodes that were chronically implanted in 1mm deep non-penetrating holes in the cranium (Woodman et al., 2007; Purcell et al., 2013; Teichert, 2016). Electrodes were positioned across the entire accessible part of the cranium at positions approximately homolog (Li and Teichert, 2020) to the international 10-20 system in the human (Li and Teichert, 2020). More details of the EEG recording setup have been provided in earlier work (Teichert, 2016; Teichert et al., 2016). Data was recorded with an Intan RHD 2000 digital amplifier. The midline electrode immediately anterior to Oz served as the recording reference and ground electrode. Data was referenced offline to the Oz electrode. In one animal, all electrodes were functional, allowing us to perform the deconvolution for all electrodes and thus visualize topographies of the F0 responses. In the second animal only a subset of the electrodes were functional, thus preventing topographical analyses.

### Pre-processing

The raw data were high pass filtered using a second-order zero-phase shift Butterworth bandpass filter with cutoff frequencies of 60 and 2000 Hz. Time-locked epochs were extracted and down-sampled to a rate of 10kHz. Epochs that exceeded an artifact-rejection criterion based on the distribution of peak-to-peak amplitudes for each individual channel were excluded from further analyses for that channel. If an epoch exceeded the relative amplitude criterion in two or more channels, it was rejected for all channels. This relative amplitude criterion allowed us to process a range of channels with different noise levels simultaneously, i.e., using the same (relative) criterion. The valid epochs were averaged separately for the four tones to obtain a total of four FFR waveforms. In addition, the valid epochs were also averaged separately for all tones and polarity to obtain eight FFRs.

### Deconvolution approach

#### Click trains

The starting point for the click train deconvolution approach were click onset times and their amplitudes. The amplitudes were further normalized to an average value of 1 across all 4 click trains. The onset times were then shifted in steps of 0.1 ms (i.e., the sampling rate of the FFRs), to create 800 identical regressors, shifted in time relative to the original F0 onsets in a range between 0 to 79.9 ms. We then fit a linear model to the FFR using all 800 regressors. To that aim, the FFRs from all stimuli and the corresponding regressors were concatenated into a single time-series padded with NaN values (**N**ot **a N**umber) between them to avoid cross-talk between the end of one stimulus and the beginning of the next. The FFR kernel was then defined as the 800 weights of the 800 regressors. The deconvolution approach thus identified the kernel, that best explained the observed FFRs as the linear sum of overlapping responses to each individual click in the click train. Note that the FFRs to all stimuli was explained by a single 80 ms long kernel. The deconvolution approach was implemented in the statistical software R, using an in-house written deconvolution package (deconvolvR).

#### Mandarin tones

An almost identical procedure was used to create the predictors for the tone FFRs. However, to create the click trains, we had placed individual clicks at the time of the peak pressure of each F0 cycle (**Fig 1**C, second F0 cycle). This choice may have been suboptimal, as peak pressure does not coincide with the timing of the actual glottal pulse. We thus identified an approach and operationalized the onset of each F0 cycle as the first positive pressure peak that coincided with a peak of power in the third harmonic (**Fig 1**C, first F0 cycle). The two different approaches yielded highly similar timing, but the estimated F0 onsets preceded the time of peak pressure very reliably by 1.01 ms. Tone FFR kernels were estimated from both types of predictors based on the timing of the peak pressure and glottal pulse. Both yielded almost identical results. However, the kernels from the peak pressure were delayed by approximately 1ms, and they explained a somewhat lower amount of variance. Furthermore, the timing of the tone kernel based on the glottal pulse matched the timing of the click kernel much better than the tone kernel based on peak pressure. Following the theoretical arguments and the empirical support, we report the tone kernels using the glottal onset time rather than the time of peak pressure.

### Quantification of model fit

The primary variable used to quantify the quality of the model fit was percent variance explained. Percent variance explained is typically calculated as 100 * (TMS-RMS)/TMS. Here RMS stands for the mean of the squares of the residuals and TMS for the mean of squares of the total signal, i.e., including variance pertaining to the actual FFR as well as measurement noise. Since no model can be expected to account for measurement noise this traditional metric cannot reach 100% unless there is no measurement noise. The limit of percent variance a model can explain is given by 100 – 100/signal-to-noise ratio. As a result, the metric is only comparable for data sets with similar signal-to-noise ratio. Because some of our data sets have rather different signal-to-noise ratios, we decided to use an alternative metric that adjusts for different SNRs. This metric sets out to quantify how much of the ‘explainable’ variance, i.e., the portion of the variance that exceeds the variance of the baseline, can be explained by the model: 100 * (TMS-RMS)/(TMS-BMS). In this context, BMS stands for the mean of the squares of the signal on the baseline, defined as the 50 ms period before stimulus onset, and the period from 320 to 390 ms after stimulus onset, i.e., 70 to 140 ms after stimulus offset. We had found the variance on the post-stimulus baseline to be systematically smaller than on the pre-stimulus baseline. Hence the decision to use the average of both periods.

Unless mentioned otherwise, we will refer to this SNR-corrected measure of percent variance explained throughout the manuscript. Percent variance explained was calculated across the entire simulation period (0 to 280 ms after stimulus onset), as well as the sustained period which excluded both on- and offset responses (50 to 250 ms). Note that in all cases, the kernel was estimated by fitting it to the entire temporal duration of the data. Consequently, any difference in percent variance explained is not caused by requiring the model to fit a simpler subset of the data, but rather depends on how well the same underlying model accounts for the data in different epochs.

Furthermore, we performed a wavelet decomposition of the signal as well as the residuals and evaluated percent variance explained in three different frequency bands, the frequency range of the fundamental frequency F0 (70–170 Hz), the frequency range of the first harmonic H1 (180–300Hz), and the frequency range of harmonics beyond the second harmonic Hx (400–1200Hz). To account for the temporal smearing of the wavelet decomposition, the time ranges of all periods were shrunk by 20 ms on each side.

#### Data split control

To prevent overfitting caused by determining the kernel and the percent variance explained from the same data set, we randomly split the data of each recording session in two equally sized subsets. The first subset of data (training set) was used to estimate the kernel. This kernel was then used to determine percent variance explained of the second subset (testing set). In the context of the work presented here, the approach was only used for the data averaged across all sessions.

#### Cross-day control

At the single session level, we used a different approach to prevent overfitting. Specifically, to explain FFRs from one recording sessions we only used kernels extracted from different recording sessions. The data fit metric for the session in question, e.g., percent variance explained, was then defined as the average of that metric using kernels from all other sessions.

#### Shuffle control

To control for the large number of predictors in the linear model (80 [ms] × 10 [samples per ms] = 800) we included a shuffle-control. To that aim we used the same averaged data and the same predictors. However, the timing of the dirac pulses was shifted such that the timing and amplitude designed to match the F0 onsets for tone 2 were used to predict data for tone 1, the timing and amplitude designed for tone 3 were used for tone 2 and so on. This approach was used for data averaged across all recording sessions as well as for data of individual recording sessions.

### Data quality and rejection of recording sessions

For the click train stimuli we recorded a total of 27 EEG sessions (animal B: 17, animal J: 14). For the mandarin tone stimuli we recorded a total of 18 EEG sessions (animal B: 2, animal J: 18). Sessions were included into the analyses if the noise of the averaged FFRs on the baseline was below 0.008 uV^2. Data quality for animal J was variable between sessions, and approximately half of the sessions did not meet the criterion (animal J, click train sitmuli: 8/14 sessions; tone: 9/18 sessions). Data quality for animal B was consistently high. Only one of the click train sessions needed to be excluded because of noise. In addition, we excluded one of the click train sessions because the signal amplitude was less than half of the other sessions, a clear outlier given the tight distribution of values for the other sessions. In summary, we used 2/2 tone sessions and 15/17 click train sessions for animal B.

Noise amplitude on the excluded sessions were distributed bimodally: a small fraction of cases with an increase of well over 10-fold, and a larger fraction with an increase below 2-fold. Including the sessions with less than a two-fold increase did not change the main conclusions. However, it did increase variability of the results between sessions and decrease the percent variance explained by a relatively modest amount. The key takeaway from including the noisier sessions is not very unexpected: if data quality is lower, less variance can be explained.

## Results

FFRs were recorded in response to two types of stimuli: (i) 4 synthetic mandarin tones using the syllable /yi/ and click train versions of these mandarin tone stimuli. Click train stimuli were created by converting the 4 mandarin tone stimuli into series of mono-phasic clicks whose timing and amplitude matched the estimated time of onset of each F0 cycle (**Fig 1**A, see Methods for details). We report data from a total of 23 EEG recording sessions using the click train stimuli (15 sessions animal B, 8 sessions animal J) and 11 sessions using the mandarin tone stimuli (2 sessions animal B, 9 sessions animal J). Each session lasted 40 minutes and contained a total of 4000 stimuli, 500 from each type and polarity.

### Tone and click train FFRs

As expected, both types of stimuli elicited periodic FFR-like responses in both animals. **Figure 2** depicts the mandarin tone stimuli as well as the grand average FFRs in the time and time-frequency domains for both subjects. In the time-domain, we observed a wide diversity of shapes of the FFRs as F0 changed both within and between different mandarin tone stimuli. In the time-frequency domain, we observed modulation of the fundamental frequency (F0) and the first harmonic (H1) in concert with the dynamically changing fundamental frequency of the mandarin tone stimuli. **Figure 3** depicts the click train FFRs in the time and time-frequency domains. The click train FFRs were qualitatively similar, but of larger amplitude than the mandarin tone FFRs. In the time-frequency domain, we observed power above the first harmonic. Especially for animal B, there was evidence of a second harmonic (F2) in cases when F0 was low, such as for click train #3 or towards the end of click train #4. Furthermore, we often observed power beyond the second harmonic in even higher frequency bands >400 Hz. In contrast to the first and second harmonic, the frequency of these higher-frequency components did not change in line with the fundamental frequency of the stimulus. These higher frequencies were also present for the tone FFRs, but harder to distinguish due to their lower amplitude. Based on the time-frequency decomposition of the FFRs, we will focus on three different frequency bands, the frequency range of the fundamental frequency F0 (70–170 Hz), the frequency range of the first harmonic H1 (180–300Hz), and the frequency range beyond the second harmonic Hx (400–1200Hz).

**Figure 2.**
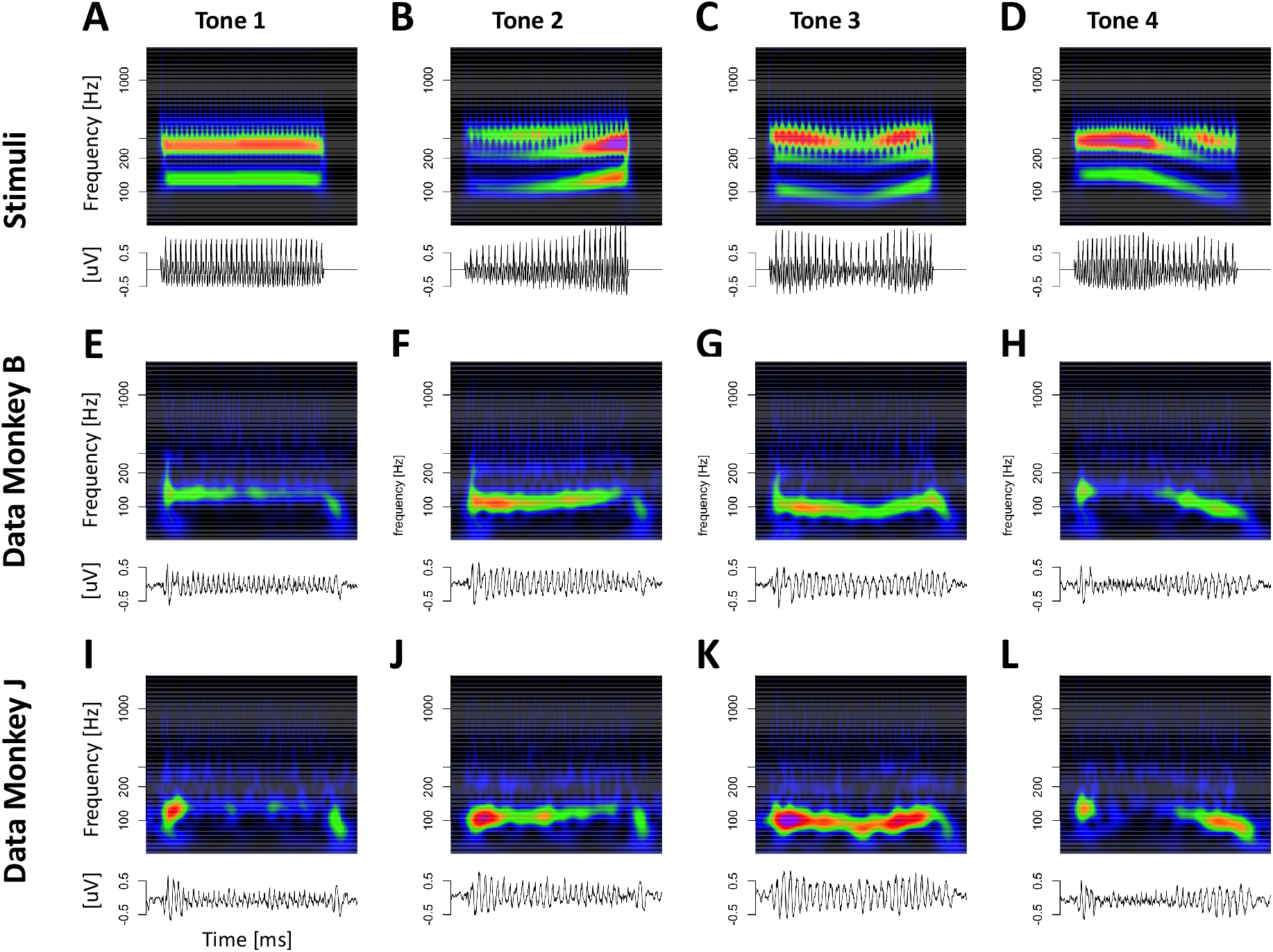
Mandarin Tone FFRs. Representation of mandarin tone stimuli and the corresponding FFRs in the time and time-frequency domain. A-D stimuli, E-F monkey B FFRs, I-L monkey J FFRs

**Figure 3.**
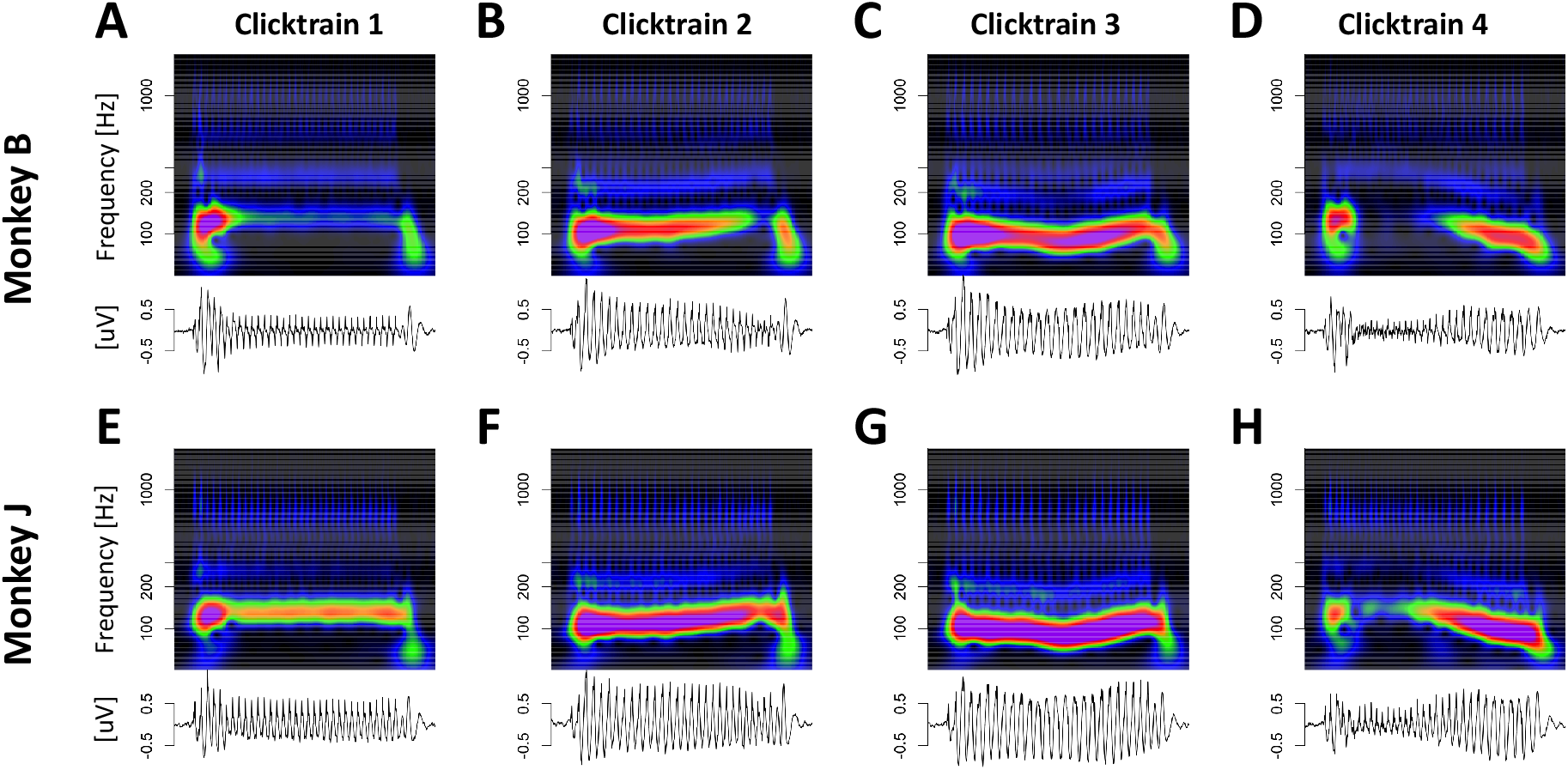
Click train FFRs. Representation of click train FFRs in the time and time-frequency domain. A-D monkey B click train FFRs, E-H monkey J clicktrian FFRs

### Deconvolution of click train FFRs

We next set out to test if FFRs with such a complex phenomenology both in the time and time-frequency domain can be explained by a simple linear super-position model. Given their larger amplitude and thus higher signal-to-noise ratio we first focused on the click train FFRs. To further improve signal-to-noise ratio, we initially focused on data averaged across all recording sessions. To that aim, data from each session was randomly split into two equally sized sets, subsequently referred to as the training and test set, respectively. Within each set, trials were averaged across the four different click train stimuli. The deconvolution was performed on the four click train FFRs averaged across all training sets. The model fit was then evaluated by comparing the model predictions derived from the training set with the data from the testing set.

**Figure 4** visualizes the deconvolution process, the F0 response, also referred to as the FFR kernel, and the model fits in the time-domain for animal B. All key features of the click train FFRs were well-captured by the convolution model (black lines in **Fig 2**C&D). It is noteworthy that the wide range of shapes of the click train FFRs could be accounted for with just one underlying kernel. The different shapes of the click train FFRs were created exclusively by slight variations of constructive and destructive interference driven by subtle timing and amplitude differences from otherwise identical F0 responses to individual clicks. In both animals, the extracted kernels contained two key spectro-temporal features: a series of brisk peaks and troughs with short latencies and high frequency, as well as wavelet-like responses at longer latencies and a lower frequency (**Fig 4**B).

**Figure 4.**
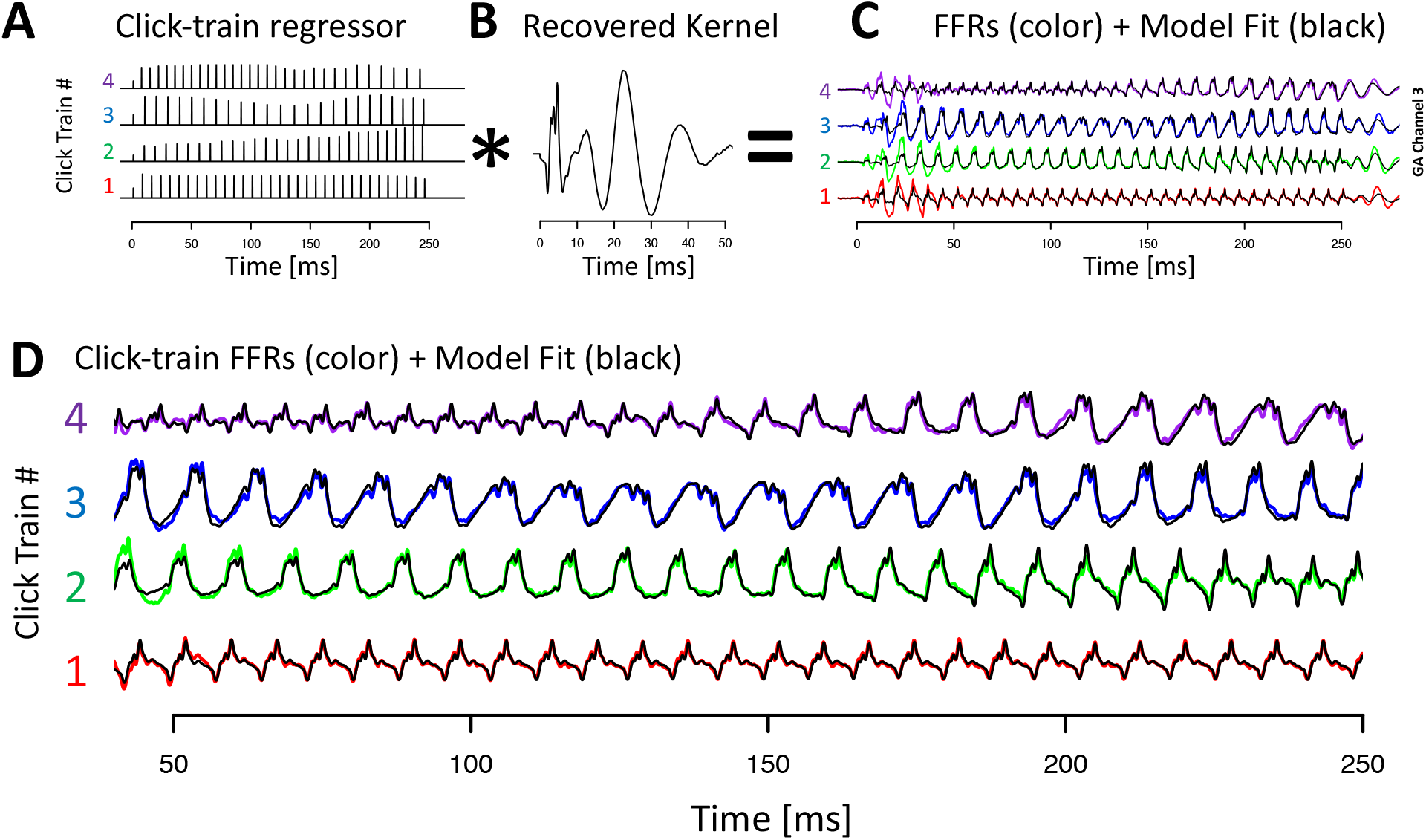
Deconvolution of grand average click train FFRs for animal J. A) Click train regressor for the four click train stimuli. The F0 contour of click train #1 matches the high tone, #2 the rising tone, #3 the dipping tone and #4 the falling tone. B) Recovered kernel which can be viewed as the impulse response to one click. C) Observed click train FFRs (color) and model fit (black). D) Enlargement of the steady-state period of the FFR response.

**Figure 5** visualizes the deconvolution process for animal J in the time and time-frequency domain. This visualization confirmed that the model captured key aspects in all relevant frequency bands and not just the fundamental frequency. Note that the model captured the components whose frequency changed dynamically with F0 (fundamental and first harmonic), as well as the higher frequency components above F2 whose frequency is unaffected by dynamic F0 of the stimulus (or the ensuing FFR).

**Figure 5.**
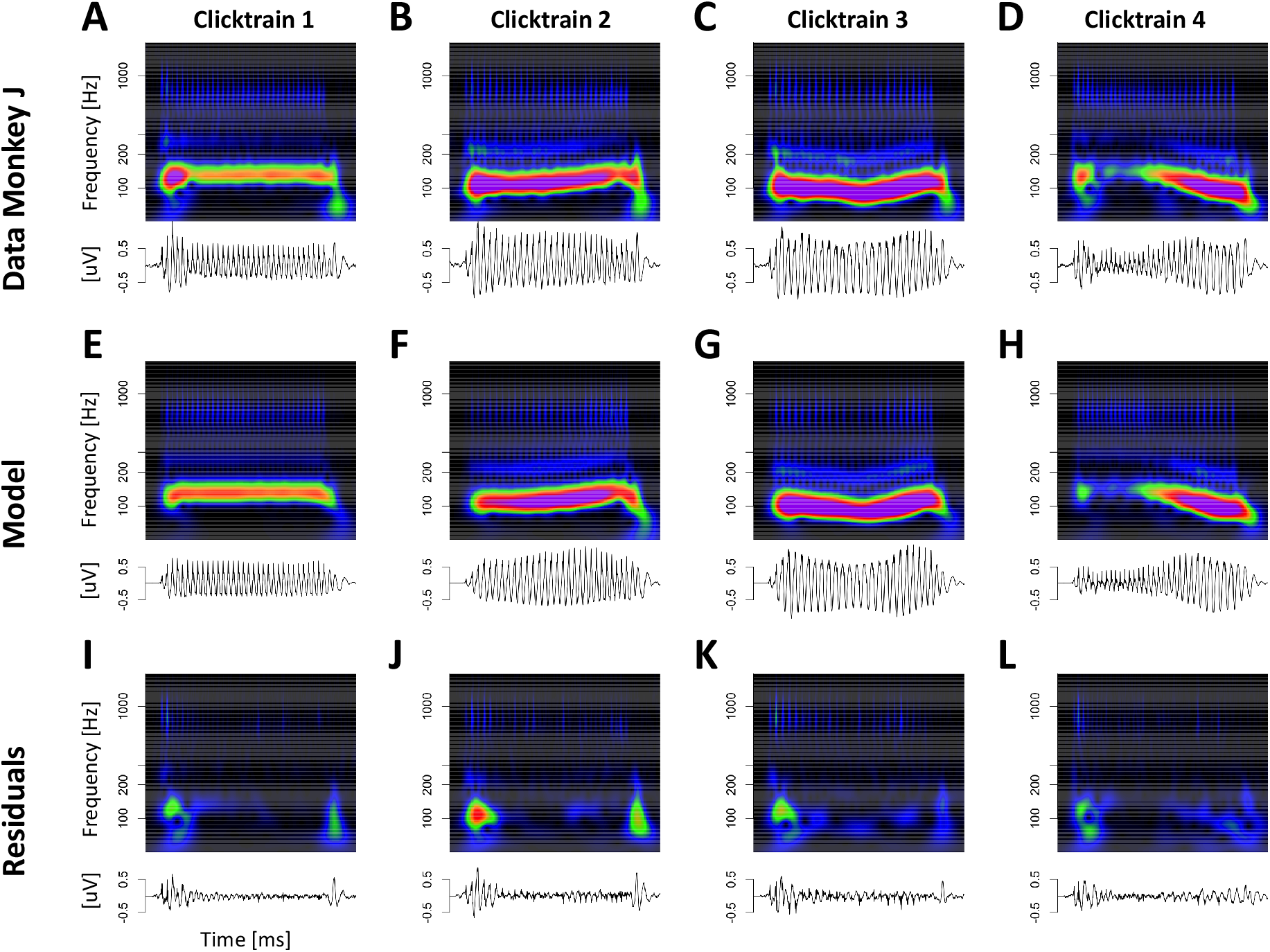
Deconvolution of grand average click train FFRs for animal J in the time and time-frequency domain. A-D) click train FFRs; E-H) fit of the deconvolution model. I-J) residuals of the model fit.

**Figure 6** visualizes the deconvolution process for the mandarin tone stimuli in the time-domain. Other than using mandarin tone FFRs as inputs to the model, the procedure for obtaining the response kernels was identical, and the results closely resembled the deconvolution process for the click train stimuli.

**Figure 6.**
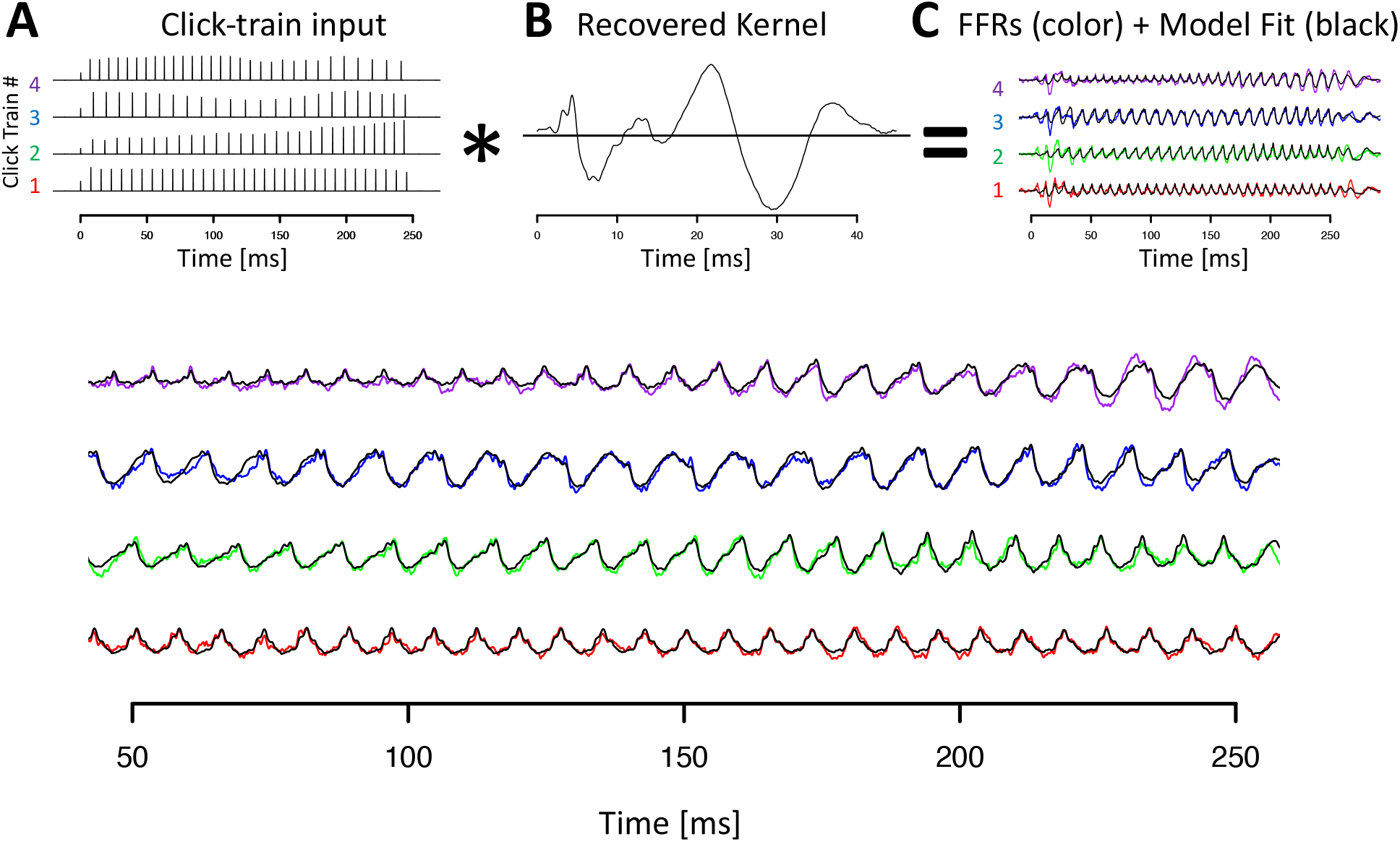
Deconvolution of grand average tone FFRs for animal J. Conventions as in Figure 4.

While the convolution model captured all key aspects of the data, we also observed regions of systematic misfit. In particular, the model underestimated the response amplitudes during the first ~50 ms of the stimulus. In part to compensate for this, the model tended to overe-stimate the amplitudes for the remainder of the stimulus. This effect may likely be caused by short-term adaptation, a non-linear effect that cannot be accounted for by a strictly linear model. We will briefly touch on this issue later in the manuscript by introducing a non-linear–linear convolution approach that resolves most of the remaining systematic misfit during the onset period.

### Percent variance explained – click train FFRs

We next quantified the performance of the model as the percent variance explained, either calculated across the entire stimulation period (0 to 280 ms after stimulus onset), or the sustained period which excluded both on- and offset responses (50 to 250 ms). Furthermore, we evaluated percent variance explained in three different frequency bands, the frequency range of the fundamental frequency F0 (70–170 Hz), the frequency range of the first harmonic H1 (180–300Hz), and the frequency range of harmonics beyond the second harmonic Hx (400–1200Hz). See methods for details.

Because no model can be expected to account for measurement noise, percent variance explained cannot exceed a threshold of 100 – 100/SNR. As a result, the traditional metric of percent variance explained is only comparable for data sets with similar signal-to-noise ratio. Thus, we decided to quantify how much of the ‘explainable’ variance, i.e., the portion of the variance that exceeds the variance of the baseline, can be explained by the model. See methods for details.

In both animals, the convolution model explained the vast majority of the explainable variance (monkey B: 79%; monkey J: 90%, solid circles in **Figure 7**A). This value was even higher in the sustained period that excluded on- and offset responses (monkey B: 95%; monkey J: 97%; solid circles in **Figure 7**B). Within the sustained period, there was a gradient of percent variance explained by frequency range. The largest fraction of variance could be explained in the F0-range, followed by the H1 and Hx ranges (F0-range: 95% and 98%, for monkey B and J, respectively; H1-range: 96% and 95%; Hx range: 92% and 92%, solid circles and lines in **Figure 7**C).

**Figure 7.**
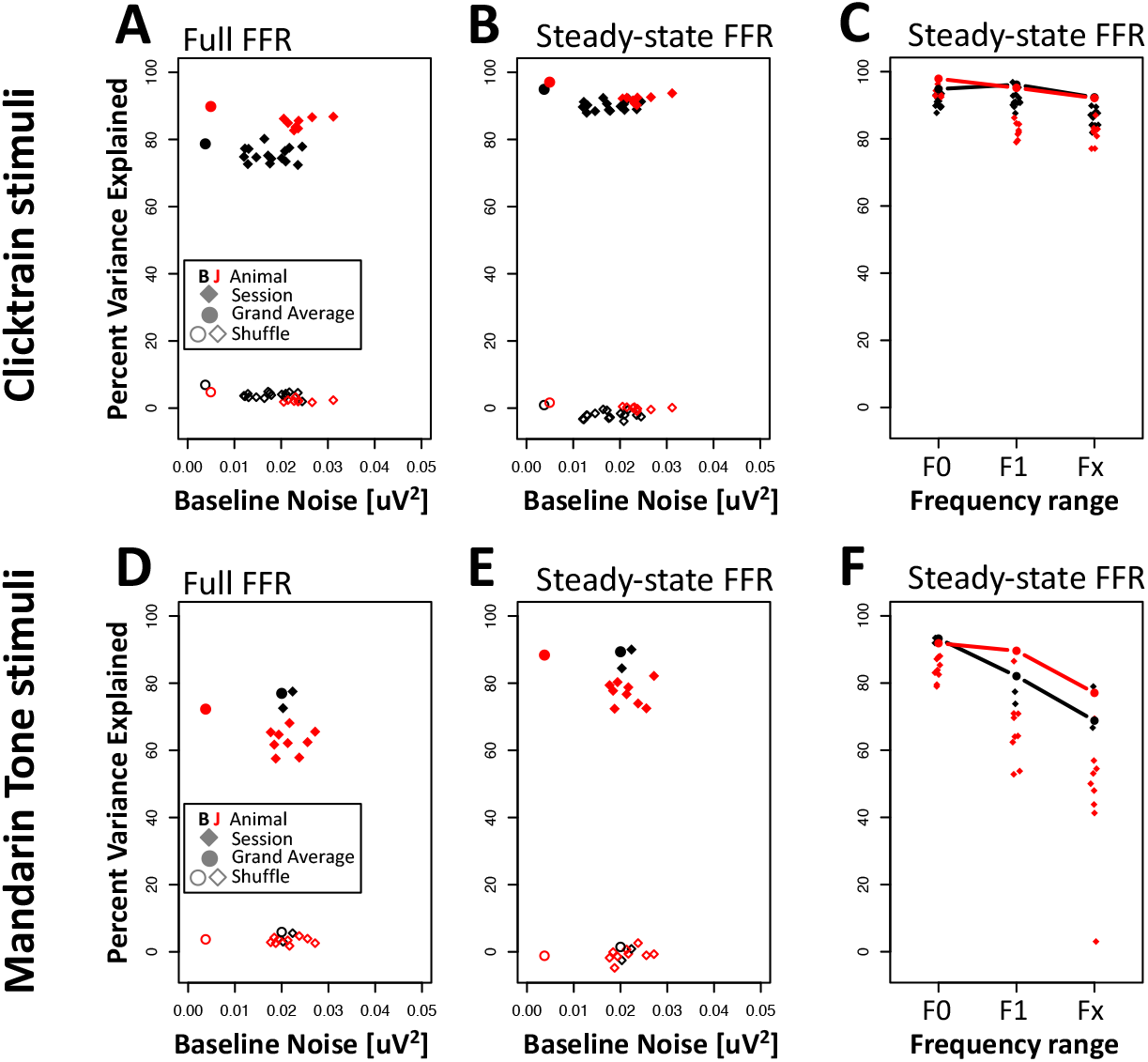
Percent Variance Explained. A) Percent variance explained across the entire FFR as a function of baseline noise. Solid points indicate fits to the grand averages across all sessions. Solid diamonds indicate fits to individual sessions. Unfilled symbols indicate fits using shuffled predictors. B) Same as A) but percent variance explained is only evaluated for steady state portion of the FFR (50-250ms). C) Percent variance explained by frequency band. E,F,G) Same as A,B,C but for mandarin tone stimuli.

We next tested if the high percentage of explained variance may be caused by overfitting. To that aim, we used a shuffle control in which the number of predictors remained constant but no longer matched the timing and amplitude of the actual F0 onsets (see methods for details). This shuffling dramatically attenuated the percent variance explained (animal B: 7%, animal J: 5%, open circles in **Figure 7**A). The percent variance explained was even smaller in the sustained period (animal B: 1%, animal J: 2%, open circles in **Figure 7**B). The lower values for the sustained period likely occurred because the shuffled model tended to capture variance at stimulus onset (which is identical for all stimuli) at the expense of the sustained period.

We next set out to quantify how much of the click train FFRs can be explained by the linear kernel in more common experimental settings, i.e., from data collected in individual recording sessions. To that aim, we calculated the kernel from data averaged across one recording session and evaluated the fit by comparing the predictions to the FFRs of all other recording sessions. The results largely replicated the findings at the level of the grand averages and confirmed that a substantial amount of the explainable variance could be captured by the linear model even at the level of individual recording sessions (animal B: 75±2.7%, animal J: 85±2.7%, mean standard deviation, solid diamonds in **Figure 7**A). An even higher percent of the variance was captured during the sustained period (animal B: 90±4.0%, animal J: 92±2.9%, solid diamonds in **Figure 7**B). Results from the shuffle control predictor confirmed that overfitting was also not a major concern for the single session data (animal B: 4±0.9%, animal J: 2±0.9%, open diamonds in **Figure 7**A). The percent variance explained by the shuffle predictor was even smaller in the sustained period (animal B: −2±1.3%, animal J: 0±0.9%, open diamonds in **Figure 7**B). The negative values for animal B indicate that the shuffle predictor inflated the variance in the sustained period.

Furthermore, the single-session analysis confirmed that the model captured the most variance in the frequency range of the F0 (animal B: 91±4.4%, animal J: 94±3.3%, solid diamonds in **Figure 7**C), followed by the frequency range of the H1 (animal B: 92±2.7%, animal J: 82±4.8%), and the highest frequency range Hx (animal B: 86±2.9%, animal J: 82±4.8%).

### Percent variance explained – mandarin tone FFRs

The results so far suggest that the deconvolution method works rather well on artificial click train stimuli. By itself, this is an important finding. However, given the substantial differences between click trains and speech, we then tested if the method also explains much of the variance of the FFRs in response to the spectro-temporally complex and realistic mandarin tones.

As for the click train stimuli, we first computed the deconvolution on data combined across all recording sessions for each animal. Kernels were fit to a training set and the quality of the fits were then evaluated by comparing the predictions to the FFRs of the test set. In both animals, the convolution model explained a large proportion of the explainable variance (monkey B: 77%; monkey J: 72%, solid circles in **Figure 7**D). This value was even higher in the sustained period that excluded on- and offset responses (monkey B: 89%; monkey J: 88%, solid circles in **Figure 7**E). Within the sustained period, there was a clear gradient of percent variance explained by frequency range. The largest fraction of variance could be explained in the F0-range, followed by the H1 and Hx ranges (F0-range: 93% and 92%, for monkey B and J, respectively; H1-range: 82% and 90%; Hx range: 69% and 77%, solid circles and lines in **Figure 7**F).

As for the click train stimuli, using the shuffled predictor dramatically attenuated the percent variance explained (animal B: 6%, animal J: 4%, open circles in **Figure 7**D). The percent variance explained was even smaller in the sustained period (animal B: 1%, animal J: - 1%, open circles in **Figure 7**E).

Despite the overall lower signal amplitudes for the tone FFRs, a large proportion of the variance was captured by the linear convolution model even on a session-by-session basis (animal B: 75±3.5%, animal J: 63±4.0%, mean ± standard deviation, solid diamonds in **Figure 7**D). Excluding on- and offset responses, the percentage variance explained is even higher (animal B: 87±3.9%, animal J: 77±4.6%, filled diamonds in **Figure 7**E). As for the grand averages, shuffling dramatically attenuated the percent variance explained at the single session level (animal B: 4.0±1.8%, animal J: 3.0±1.9%, open diamonds in **Figure 7**D; sustained period: animal B: −1 ±2.4%, animal J: −1 ±2.9%, open diamonds in **Figure 7**E), again confirming that overfitting was not a substantial contribution to the high percentage of variance explained.

Furthermore, the single-session analysis confirmed that the model captured the most variance in the frequency range of the F0 (animal B: 93±1.1%, animal J: 84±3.8%, solid diamonds in **Figure 7**F), followed by the frequency range of the H1 (animal B: 76±2.5%, animal J: 66±11.1%), and the highest frequency range Hx (animal B: 73±8.7%, animal J: 47±19.5%).

### Consistency of deconvolution approach across recording sessions

The ability to explain the FFRs of one recording day using the kernel from a different session, suggests that the kernels are remarkably similar between days. **Figure 8** A&B confirms the high degree of similarity for the click train kernels. Especially early features of the kernel (<5 ms) were highly preserved across sessions, to the point that it was hard to even distinguish the presence of more than one trace. Above 5 ms, differences between sessions became somewhat more apparent. The largest between-session variability was observed for the late wavelet-like response between 15 and 35 ms. We quantified the similarity of the kernels as the Pearson correlation coefficient which was found to be 0.97±0.02 for both animals (average plus minus standard deviation). Note that while the kernels for different sessions were highly similar, the kernels for the two animals were quite distinct from each other. In particular, the early features of the kernels below 5 ms are like a finger-print that uniquely identify the subject with high confidence on the basis of a single session.

**Figure 8.**
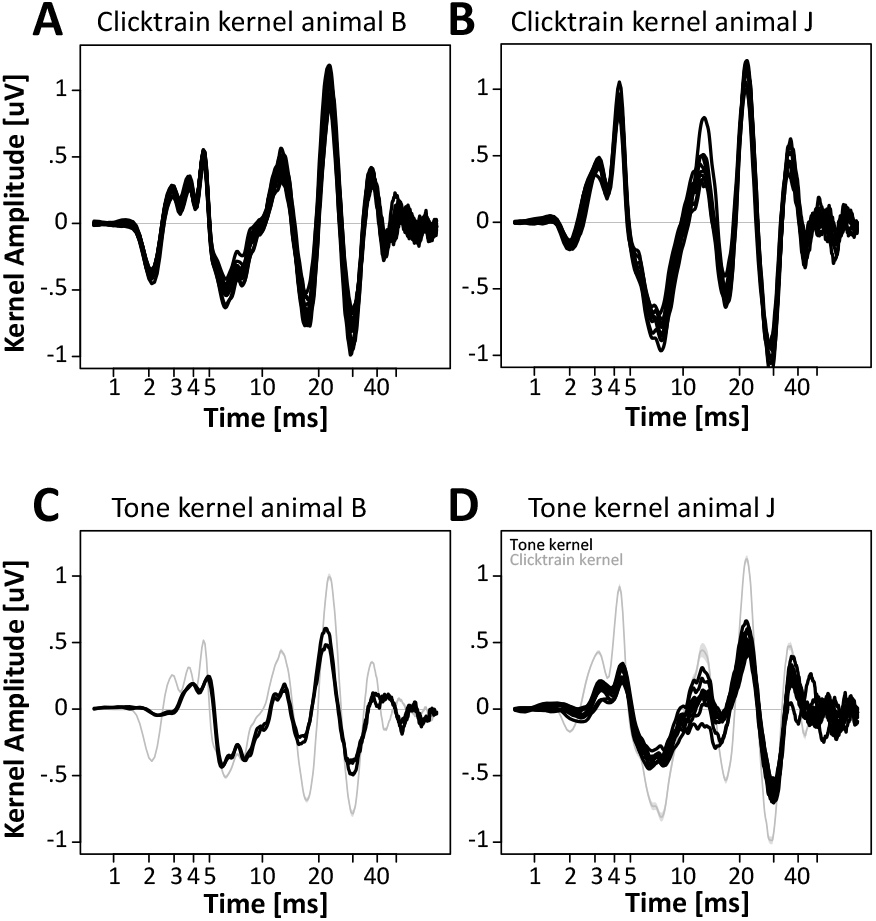
Comparison of F0 responses across sessions, stimuli and subjects. A,B) Click train F0 responses for individual sessions of animals B and J. C,D) Mandarin tone F0 responses for individual sessions of animals B and J.

Cross correlations for kernels of the mandarin tone stimuli were similarly high (animal B: 0.98±NA, animal J: 0.91 ±0.08). Standard deviation was not available for animal B, since only two sessions were recorded, resulting in a single cross-correlation value. For monkey J, the average cross-correlation was attenuated mostly by one session. As a result of the left-ward skew of the distribution, the median correlation coefficient was a good bit higher and probably a more robust estimate (median correlation coefficient monkey J: 0.95).

### Spectro-temporal features of the F0 responses

**Figures 8**&**9** visualize the time-course of the click train kernels for both animals. The kernels could be split into three epochs: (1) an initial period from 1-5 ms that featured a series of brisk peaks and troughs; (2) a transition period from 5 to 15 ms; (3) the final period from 15 to ~45 ms that featured 3 peaks and 2 troughs of a large-amplitude and relatively slow, wavelet-like oscillation. In the initial period, both animals exhibited a prominent trough at ~2 ms and a prominent peak at ~4.5 ms. In between the two, animal B featured two peaks at 2.9 and 3.7 ms, while animal J featured only one intermittent peak at 3.1ms. The peak at ~4.5 ms likely corresponds to wave V of the brainstem auditory evoked potential. Transforming the kernels into the time-frequency domain revealed a complex spectral composition (**Figure 9**A&B top panels). Both animals exhibited prominent high-frequency components above 500 Hz: in animal B, they manifested in two distinct spectral peaks at 600 and 1050 Hz. In animal J, they manifested as a single peak at 700 Hz. In both animals, these high-frequency bursts were extremely short-lived. In addition, both animals show spectral power at frequencies around 200 Hz. For both animals, activity in this frequency range extended into the transition period. The key spectro-temporal feature of the kernel was an extended period of power in the lower frequency range between 70 and 120 Hz. Closer inspection revealed a gradual decrease of frequency over time: in animal B the frequency decreased from 90 Hz to 70 Hz, in animal J the frequency decreased from 105 to 75 Hz. It is unclear if this decrease resulted from the gradual change of frequency of a single component, or from the transition between two components with slightly different frequencies.

**Figure 9.**
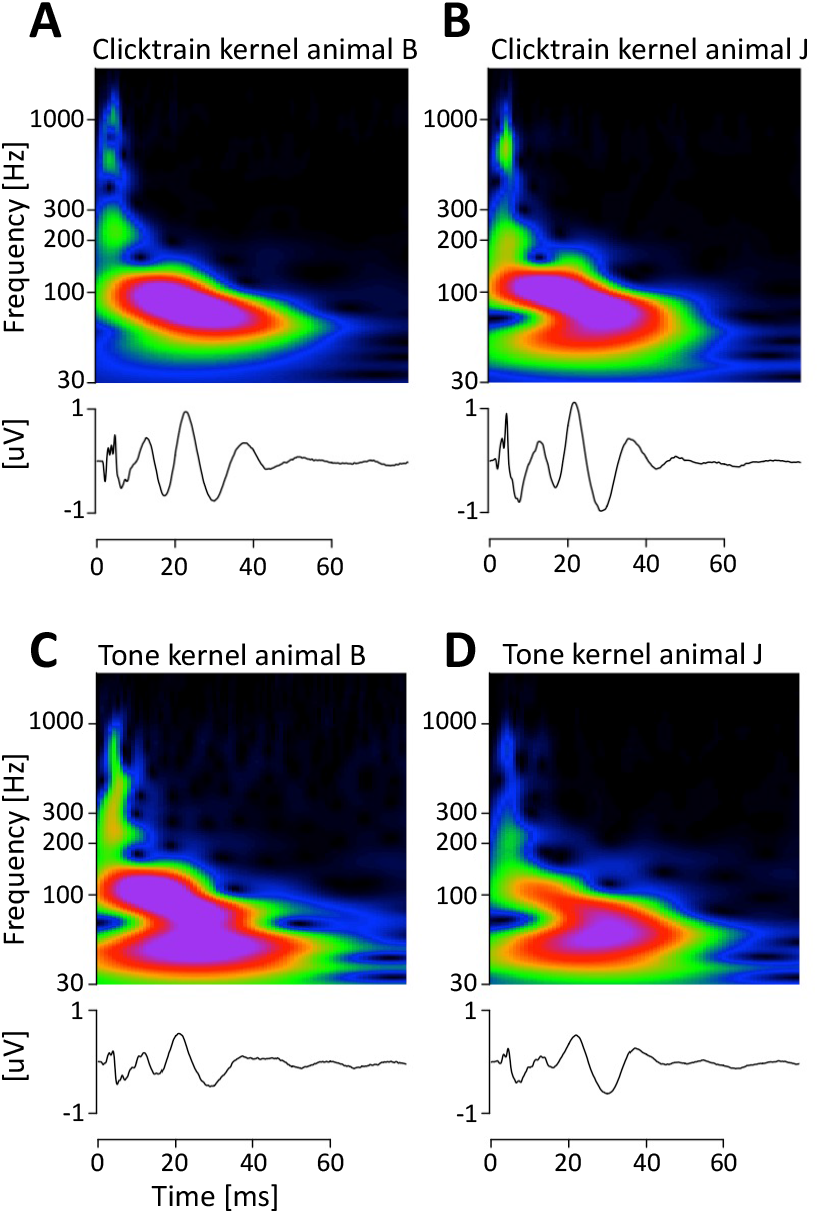
Comparison of F0 responses across stimuli and subjects in the time-frequency domain. A,B) Average click train F0 responses for animals B and J. C,D) Average mandarin tone F0 responses for animals B and J.

**Figures 8**C&D provides a direct comparison of the tone and click train kernels in the time domain. The most striking difference is the overall reduced amplitude of the kernels, which matches the overall reduced amplitudes of the tone FFRs themselves. However, most of the key features of the kernels were preserved. In particular, the timing and polarity of most peaks were identical. Only the earliest putative brainstem components were affected more strongly. In both animals the initial trough that was evident at ~2 ms for the click train kernels was reduced in amplitude, temporally smeared and delayed to ~3 ms. In animal B, this temporal smearing may have contributed to the cancellation of the first of the three subsequent positive peaks that occurs at 2.9 ms in the click train kernel. **Figure 9** highlights another interesting distinction that is not visible in the time-domain. For both animals, the tone kernels included power in an even lower frequency band centered around 50 Hz that was not active for the click train kernels.

### Topography of the click train F0 responses

It is tempting to link these different spectro-temporal features of the kernel to processing in brainstem, midbrain and cortex, respectively. If correct, it would support the notion that the deconvolution method was indeed able to partially disentangle these different generators whose activity is temporally completely overlapping in the FFR. If different latencies of response components in the FFR kernel indeed reflect the gradual activation of successively higher stages of auditory processing, then this should be reflected in different topographies for early relative to late components. In one subject, animal B, we had access to an entire grid of 33 EEG electrodes. We thus set out to estimate the kernels for all 33 EEG electrodes in this animal. The resulting topographies are summarized in **Figure 10**. The topographies of the putative cortical components indeed closely resembled the topographies of classical evoked potentials that are believed to arise from core auditory regions in the superior temporal plane (Teichert, 2016). In contrast, the putative brain-stem topographies were much more varied, and, except for the peak at 4.2 ms, clearly not of cortical origin. The topographies of the putative midbrain components were diverse. While the topography of the component at 6 ms was not unlike the classical cortical topography, the component at 11ms was clearly not suggestive of cortical origin.

**Figure 10.**
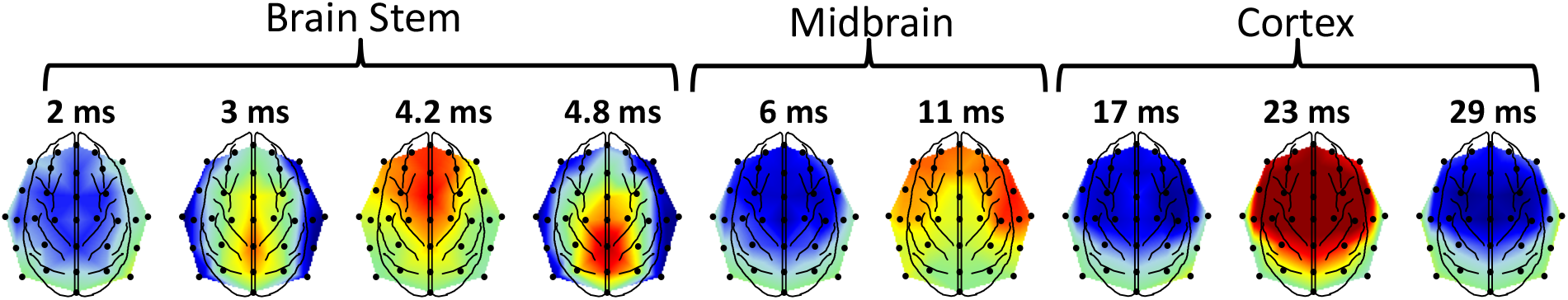
Topography of click train F0 responses. Topography of different peaks and troughs of the F0 onset response for animal B. Different components are tentatively grouped in to brainstem, midbrain and cortex based on latency, frequency and topography.

### Dependence of F0 responses on stimulus polarity

Both stimulus types, click trains and mandarin tones, were presented in two randomly intermixed polarities. The results presented above reflect the model fit to the average of the two polarities. **Figure 11** compares the FFRs for the two polarities for the click train stimuli and mandarin tone stimuli separately. For the click train stimuli, the two polarities were quantitively almost identical, except for a minor deviation during the putative midbrain components at a latency of ~7 ms. Note that while the effect was extremely small in absolute terms, it was highly replicable between sessions and present in both animals. A qualitatively similar, but substantially larger effect emerged for the tone kernel: the difference between the two polarities was most evident in the late brainstem and early midbrain latencies. In both animals, the putative component V of the brainstem response was strongly attenuated in the rarefaction condition (**Figure 11** B&D, orange arrow). In its stead, a new peak at a latency of ~7ms that was superposed over the trough was also observed at this latency (**Figure 11** B&D, blue arrow). Note that for both stimuli and both animals, the putative cortical contribution to the kernels beyond ~15-20 ms onwards are again very similar. As a result, the temporal fine-structure, i.e., the subtraction between the polarities, would emphasize the putative midbrain components that emerge at latencies between 4 and 15 ms, at the expense of subtracting out the putative cortical contribution as well as the earliest brainstem components.

**Figure 11.**
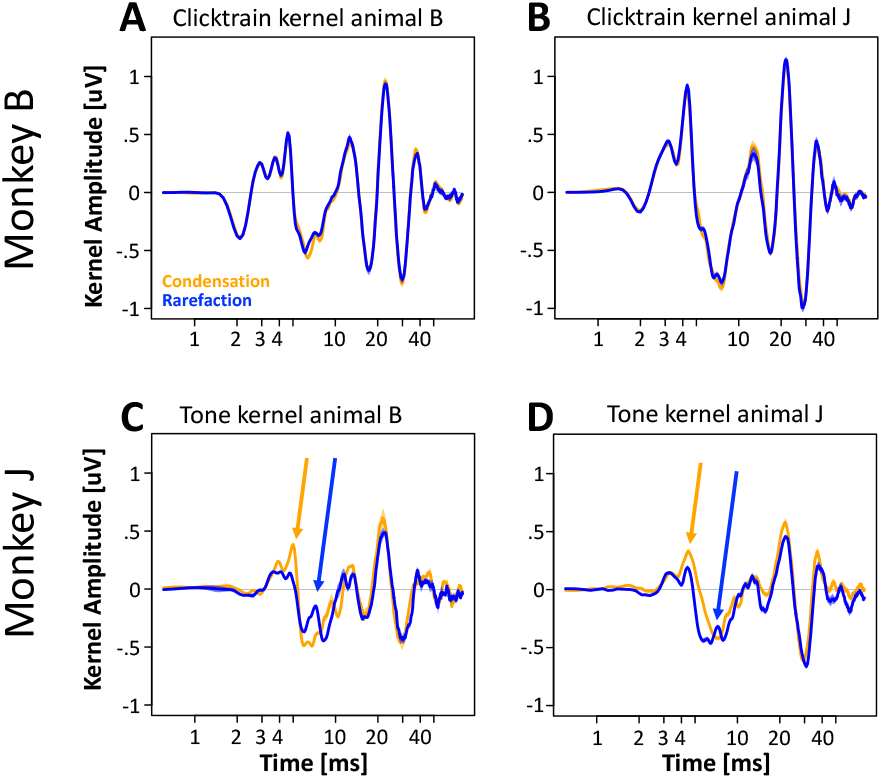
Effect of stimulus polarity on F0 responses. A,B) Effect of stimulus polarity on click train kernels for monkeys B and J (orange: condensation, blue: rarefaction). C,D) Same for mandarin tone kernels.

### Non-linear – linear deconvolution model

For both stimulus types, the linear model could predict a surprisingly large amount of the variance. However, in both cases, even the click train FFRs, the linear model fell short of explaining a substantial amount of variance around stimulus onset. The observed pattern of misfit suggests that short-term adaptation prevents the linear model from providing an even better account of the data. To confirm this hypothesis, we developed a two-stage model that includes a non-linear first stage to account for short-term adaptation, and the linear convolution model as a second stage. The short-term adaptation model uses two parameters, tau and U, to estimate how quickly and how strongly early stages of the auditory system adapt to the repeated F0 onsets. In contrast to the previous convolution model, the convolution stage of this non-linear – linear model also included a stimulus onset regressor to allow for the possibility that the very first F0 onset triggers an onset response that is qualitatively distinct from the remaining F0 responses. To keep the number of regressors similar and the gradient descent computationally manageable, we reduced the lag from 80ms (800 regressors) to 45 ms for both type of response (2 × 450 = 900 regressors). As before, the model was fit to the training set and model fit was evaluated in the testing set.

In both animals, the non-linear – linear convolution model improved model fits for the click train stimuli, especially in the onset period (monkey B: 58% to 91%, monkey J: 72% to 92%). Noticeable improvements could also be found when focusing on the entire FFR period (monkey B: 79% to 92%; monkey J: 90 to 94%). Importantly, percent variance improved or remained constant even in the sustained period (monkey B: 95% to 97%; monkey J: unchanged at 97%), even though fewer degrees of freedom were used to model the sustained period (rather than 800 parameters, the non-linear – linear model used only 2 non-linear parameters plus 450 F0 response parameters to model the sustained period; the 450 predictors for stimulus onset have no direct effect on the sustained period). Similar improvements were found for the mandarin tone stimuli in the onset period (monkey B: 55% to 90%, monkey J: 42% to 91%), across the entire FFR (monkey B: 77% to 88%, monkey J: 72% to 87%), and in the sustained period (monkey B: 89% to 92%, monkey J: 88% to 88%).

The time-constants tau of the short-term synaptic depression that provided the best fit were well below 100 ms for the click train stimuli (monkey B: 63 ms, monkey 26: y ms) and the mandarin tone stimuli (monkey B: 74 ms, monkey 13: y ms). Such short time-constants are consistent with a locus of adaptation in the early auditory system.

## Discussion

In this study, we characterized a deconvolution approach that recovered F0 responses from FFRs in response to stimuli with time-varying pitch in the non-human primate. Our ultimate goal is to link altered FFRs observed in neuropathologies to specific latencies of the F0 response and thus to narrow down the anatomical substrate of the pathologically altered FFRs. Such an approach would be particularly useful in clinical settings that often record FFRs with a simple three-electrode montage (Bidelman, 2015), and are thus not amenable to sophisticated source reconstruction analyses.

We report advances along three main avenues. First, we were able to show that the convolution model captures a substantial portion of the variance of the mandarin tone and click train FFRs. Second, we were able to show that the kernels indeed have distinct spectro-temporal features that emerge at distinct latencies and likely reflect the sequential activation of generators in brainstem, midbrain and cortex. Third, we were able to show that the FFR kernels can be estimated with high signal-to-noise ratio. In the following we will discuss the implications of these advances in more detail.

### F0 onset response captures much of the variance of mandarin tone and click train FFRs

A key novelty of our study is that it allowed us to quantify how much variance of the FFRs can be explained by the F0 responses. This is important, because it determines the likelihood that the approach will be able to account for altered FFRs in future work. To clarify why this is so important, we point out that the convolution approach can be viewed as data compression algorithm: complex and high-dimensional FFRs consisting of ~12,000 data points (4 tones times ~300 ms duration times 10 samples per ms) are represented by a much simpler kernel consisting of 800 data points (80 ms duration times 10 sample per ms). As with any data-compression algorithm, and especially for one with such a high compression ratio, its utility is determined by the amount of information loss. The less variance the algorithm captures, the more likely is a scenario where FFRs differ meaningfully between conditions but the F0 responses do not, simply because the relevant features of the FFRs were not captured by the linear model.

In the best-case scenario, i.e., when excluding on and offset responses and when using high signal-to-noise grand averages, the F0 responses can account for an average of 96% of the variance of the click train FFRs and for 88% of the variance of the tone FFRs. Even at the level of single sessions, the model was able to explain on average 91% of the variance for the click train FFRs and 82% of the variance for the tone FFRs. Our finding that such a substantial portion of the FFRs was explained by the convolution method increases the odds that F0 responses will be able to capture many clinically relevant FFR phenomena. Since the F0 responses capture more variance for the click train FFRs, one could argue in favor of using the click train stimuli in clinical settings. However, this would only be warranted if the click train FFRs can be shown to be equally sensitive to pathological changes as other commonly used FFR stimuli.

It is worth noting that the F0 responses are less adept at capturing variance in the higher frequency ranges. This drop-off is particularly pronounced for the mandarin tone stimuli and for single sessions (rather than grand averages). In those circumstances, the utility of the method will likely be reduced. However, it is worth noting that even if the F0 response captures less variance in the higher frequency ranges, it does not automatically mean that it won’t be sensitive to any neuropathological changes in that frequency range. It is quite possible for the pathological alterations to occur within the linear subspace that can be captured by the F0 response. In this case, it would even be expected to have higher sensitivity than the FFRs themselves because they would ignore anything outside of that linear subspace. However, capturing less of the variance would certainly suggest that it is more likely for at least some of the pathological changes in the higher frequency range to be outside of the linear space of the kernel.

### F0 responses compress FFRs into a meaningful format

We were also able to address a second key question that determines the utility of the deconvolution approach, namely whether or not the F0 responses represent information about the FFRs in a meaningful format. Specifically, we had speculated that the latency of different features of the F0 response would represent the latency of different neural generators being activated sequentially along the ascending auditory hierarchy. Indeed, we were able to identify distinct spectro-temporal features that emerge at distinct latencies and likely reflect the sequential activation of generators in brainstem (<5ms; 400-1000 Hz), midbrain (5-15 ms; 180-300 Hz) and cortex (15-45 ms; ~90 Hz). This hypothesis was supported by distinct topographies that could be measured in one of the two animals that had a fully intact 32 channel EEG system.

These results are consistent with and extend some closely related earlier studies. Bidelman tested if the FFR to a click train stimulus can be explained as the superposition of an empirically measured 12 ms long auditory brainstem responses to each click in the train (Bidelman, 2015). The conclusion from the paper was that the FFR was not satisfactorily explained by auditory brainstem responses, suggesting that other structures must contribute to the FFR. Our results are consistent with this conclusion. In order to explain the FFRs well, it was necessary to allow the kernel to be at least 45 ms long, thus extending well beyond the temporal range of auditory brainstem latencies. Our results are also consistent with an earlier study showing that the auditory steady-state response can be modeled as the linear superposition of onset responses to each individual 40Hz cycle (Bohórquez and Özdamar, 2008). Our findings extend this work into a higher frequency range and into the realm of spectro-temporally complex speech sounds. More recent work, conducted in parallel with studies reported here, has used a similar deconvolution approach to calculate the F0 response from continuous speech (Polonenko and Maddox, 2021). In line with our findings, they also identified F0 responses that are consistent with the notion of resulting from a sequential activation of generators along the ascending auditory pathway. Our work extends their findings by showing that F0 responses account for the bulk of the FFRs, and likely also speech-evoked responses in general. In addition, our results point out the limitations of the linear superposition approach and how to by including a simple short-term adaptation component that adjusts the effective amplitudes of the F0 cycles based on their actual amplitude as well as adaptation caused by the processing of previous F0 cycles.

For the type of F0 responses described above to arise, neural activity in the early auditory system needs to be preferentially focused on a specific phase of the F0 cycle. Furthermore, the sharp spectro-temporal features of the measured F0 responses suggest that the focusing of neural activity must have been temporally very precise. For the click train stimuli, such highly precise focusing is easily explained by the fact that all of the acoustic energy itself is focused on one specific phase. However, in the case of the mandarin tone stimuli, acoustic energy is distributed rather evenly throughout the entire F0 cycle (**Fig 1**C). Hence, it was not clear a priori if the focusing of neural activity to a specific phase would be strong enough and temporally precise enough to generate an F0 response in the same way as for the clicks in the click trains. In the context of this manuscript, we had operationalized the onset of the F0 cycle of the mandarin tones as the positive pressure peak (in the time-domain) that coincides with the first and strongest of six closely spaced peaks of power in the third formant (in the timefrequency domain) that follow the opening of the glottis. Of all spectro-termporal features, this one probably comes closest to resembling a click. Nevertheless, it is important to note that this feature only represented a very small portion of the total acoustic energy of each F0 cycle. Hence, it is somewhat surprising, that the mandarin tone F0 responses are only moderately smaller and share so many striking similarities to F0 responses of the click trains. It remains to be determined to what degree our particular choice of stimulus, i.e., the /yi/ vowel with its very high third format, or some non-linearities of the early auditory system, e.g., refractory periods or other forms of extremely short-term adaptation, facilitate this process of focusing neural responses to a specific phase of the F0 cycle.

The key difference between the tone and click train F0 responses is the absence of the first negative and the first positive ABR-like potentials at latencies of 2 and 3 ms, respectively. We speculate that their absence is caused by temporal smearing of the earliest auditory responses to spectro-temporal features distributed across the entire F0 cycle. This interpretation implies that the focusing of responses to a specific phase of the F0 cycle becomes more precise at later processing stages of the brainstem.

It remains to be determined why the F0 responses depended at least in part on the polarity of the mandarin tone stimuli. But it is noteworthy that the differences were strongest in the latency range of 5 to 10 ms that most likely reflect early processing stages in the midbrain. Interestingly, the putative cortical components of the F0 response were mostly insensitive to polarity. Taken together, these findings suggest that the temporal fine structure of the FFRs which is typically analyzed by subtracting FFRs to the different polarities would preferentially enhance neural activity originating from the midbrain, while specifically attenuating cortical responses.

### F0 responses can be measured with high signal-to-noise

Finally, we were able to show that the F0 responses can be estimated with high signal-to-noise ratio. The mean pairwise correlation coefficient between F0 responses estimated on different days was above .90 for both animals and both stimulus types. Such a high signal-to-noise ratio is possible because F0 response is estimated from approximately 120,000 F0 cycles (4000 trials, each of which contains on average 30 F0 cycles). The high signal-to-noise ratio of the F0 responses suggest that even small effects can be detected with a very reasonable number of sessions or subjects and may thus provide a solid basis for downstream statistical inference.

### Limitations of the linear convolution model

The high degree of variance that can be captured with the F0 responses suggests that the neural responses to each click in the click train were able to propagate through subsequent stages of the auditory processing hierarchy largely without interference from previous or subsequent clicks that were being processed at the same time in higher or lower processing stages. Given the rich recurrent connections between different stages of the auditory hierarchy, and numerous well-established non-linearities at the earliest stages of auditory processing (Heinz et al., 2001; Dau, 2003; Zilany et al., 2014), one might have predicted that a linear convolution model would be sorely insufficient to capture much of the spectro-temporal complexity of the FFRs.

However, it is also important to keep in mind that the linear model fell short of capturing all of the variance, especially around stimulus onset. Accounting for stimulus onset with an additional onset regressor and allowing the amplitudes of the click-responses to be subject to short-term adaptation was able to increase percent variance explained to above 90% even in the onset period. These results show that relatively minor deviations from the assumption of linearity can lead to substantial additional improvements.

### Future directions

While the results so far are promising, several additional steps need to be taken before the method can be used to identify which processing stages are the cause of altered FFRs. Most importantly, the findings need to be confirmed in humans. Our own preliminary results (Teichert et al., 2020) as well as work with continuous ‘peaky’ speech (Polonenko and Maddox, 2021) suggest very similar effects in humans. But the percent of variance that is captured by the F0 responses remains to be determined for human participants. Furthermore, it is likely that at least initially latency by itself is not sufficient to unequivocally identify an underlying generator. Even for extremely well-established classical onset responses such as the different ABR waves or the different mid-latency components, there is considerable debate about their more finegrained origin. Consequently, we predict that the method should initially be calibrated in a sample data set with high-density EEG/MEG recordings to leverage both latency and topography of the F0 responses. Once the origin of different peaks and troughs has been established, subsequent analyses will no longer be reliant on high-density EEG recordings.

Furthermore, the ability of the deconvolution approach to correctly identify generators based on the latency of the F0 responses needs to be validated empirically by recording FFRs directly from these structures. Ongoing work takes advantage of the monkey model system to acquire these invasive recordings and confirm the role of cortex to the later components of the F0 response.

It is known that the FFR fine-structure can mirror the formant structure of the underlying vowel (Arenillas-Alcón et al., 2021). The current experiments were performed exclusively using the vowel /yi/, so it remains an open question if and how the F0 responses may be modulated by the formant structure of different vowels.

The current version of the click train stimuli used clicks with twice the peak amplitude of the corresponding F0 cycle to account for the bi-directional modulation of the speech sounds. However, going forward, the amplitude of an F0 cycle may need to be defined in a more principled way. Specifically, our results suggest that the F0 responses were driven by a very specific aspect of the F0 cycle: a click-like response driven mostly by a peak of power in the third formant. Thus, it may be more appropriate to define the amplitude of the regressor as the amplitude of this specific aspect of the F0 cycle.

### Conclusion

Based on our studies in the rhesus macaque, we conclude that the deconvolution method can be used to compress complex and high-dimensional FFRs to stimuli with timevarying pitch into a short and meaningful F0 response that captures most of the variance of the FFRs. The different latencies of the peaks and troughs likely reflect the sequential activation of structures along the auditory pathway, and may at some point be useful to map altered FFRs in disease to altered function in specific brain regions.

There are already a large number of different ways to analyze FFRs, including broadband timing, F0 periodicity, phase consistency, and stimulus response correlation, to name just a few (Krizman and Kraus, 2019). The deconvolution approach and the resulting F0 responses are thus just one of many currently available analysis tools. We propose that the strength of the deconvolution approach arises from the three main points outlined above: 1) the F0 responses are a lower-dimensional summary that captures and condenses much of the variance of the original FFRs, 2) the latency of different features of the F0 responses is meaningful, and likely reflects the latency of different generators, thus linking altered F0 responses to specific anatomical substrates, and 3) the F0 responses can be measured with high signal-to-noise ratio, thus increasing the sensitivity and power of subsequent statistical analyses.

